## supplemental file 1 for "Unveiling the Domain-Specific and RAS Isoform-Specific Details of BRAF Regulation"

**Supplementary File 1**

**Unveiling the Domain-Specific and RAS Isoform-Specific Details of BRAF Regulation**

Tarah Trebino^1^, Borna Markusic^1,2^, Haihan Nan^1,3^, Shrhea Banerjee^1^, Zhihong Wang^1,4^

^1^Rowan University, 201 Mullica Hill Rd, Glassboro, NJ 08028

^2^Max Planck Institute of Biophysics, Max-von-Laue Straße 3, 60438 Frankfurt am Main, Germany

^3^School of Laboratory Medicine and Life Science, Wenzhou Medical University, Wenzhou, Zhejiang, China 325035

**NT2 +/- HRAS HDX peptide plots**

Plots are numbered from the start of the 6xHis/MBP tag as residue 1. BRAF starts at residue 409. Blue= NT2-apo; Magenta= NT2+HRAS. BRAF residues 1-152 (BSR) = peptides 409-560. BRAF residues 153-224 (RBD) = peptides 561-632. Cyan boxes highlight increased H-D exchange rate; Red boxes highlight decreased H-D exchange rate.

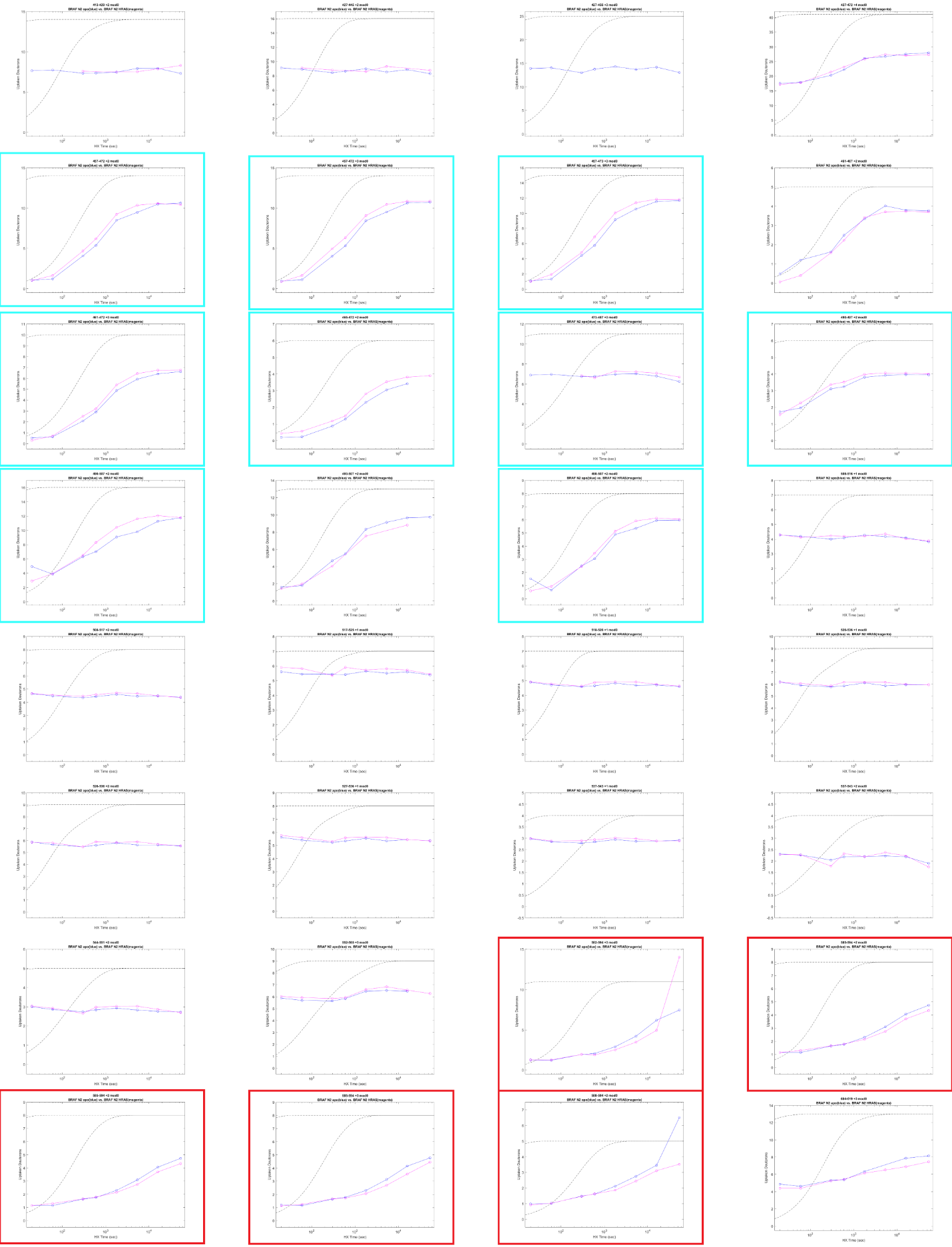

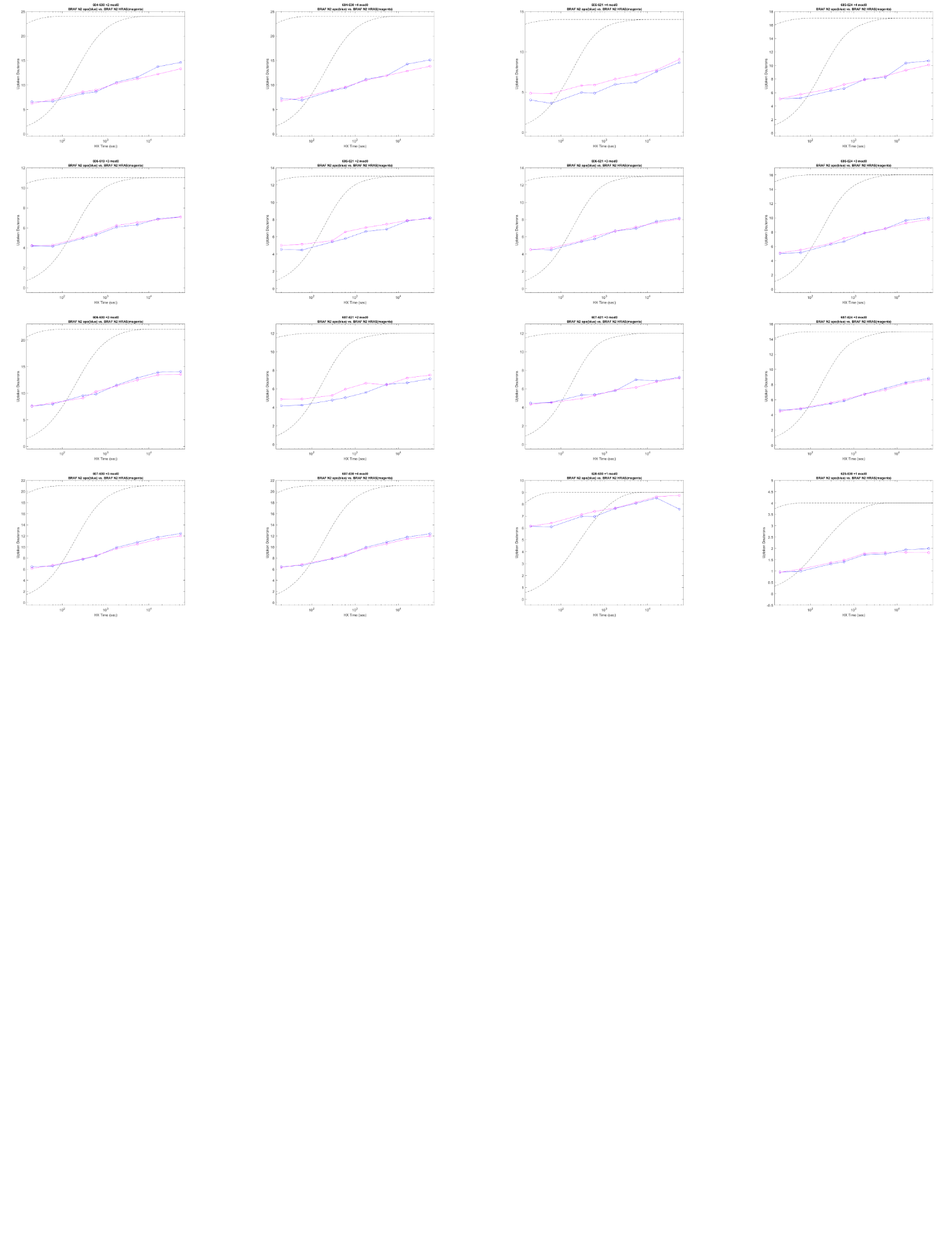

**NT3 +/- HRAS HDX peptide plots**

Plots are numbered from the start of the 6xHis/MBP tag as residue 1. BRAF starts at residue 408. Blue= NT3-apo; Magenta= NT3+HRAS. BRAF residues 153-224 (RBD) = peptide residues 412-483. BRAF residues 232-284 (CRD) = peptide residues 491-543. Red boxes highlight decreased H-D exchange rate.
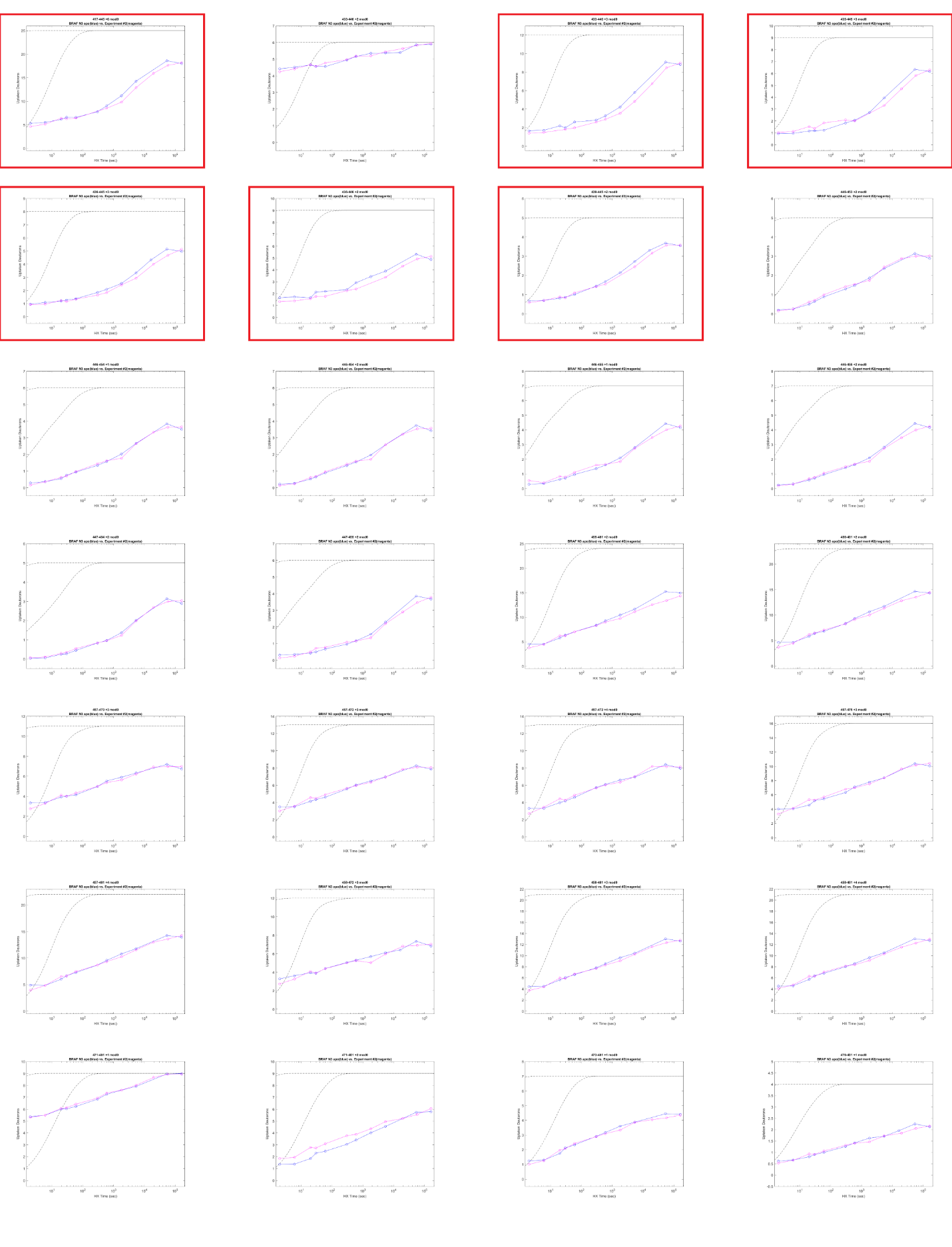

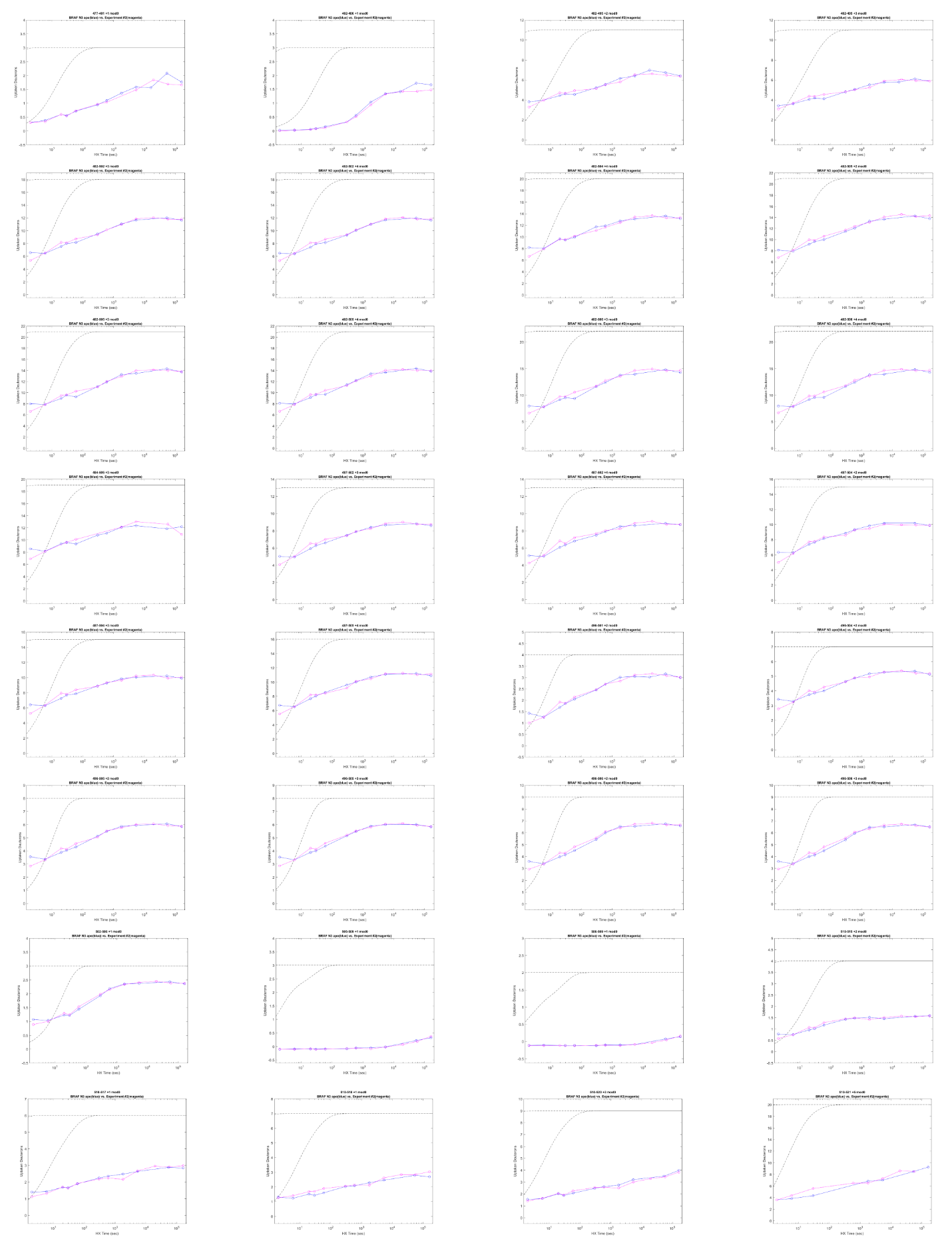

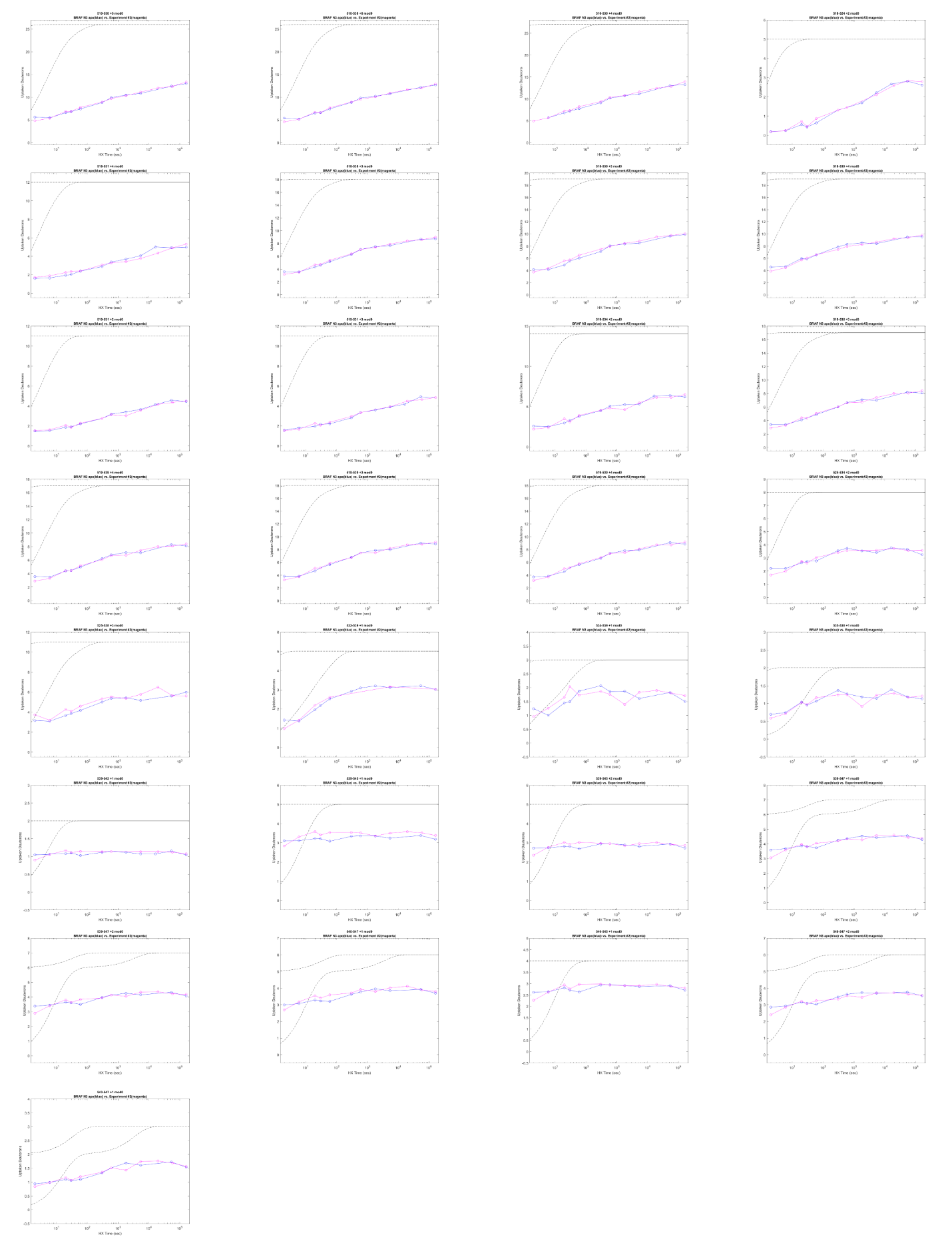

**NT2 +/- KRAS HDX peptide plots**

Plots are numbered from the start of the 6xHis/MBP tag as residue 1. BRAF starts at residue 409. Blue= NT2-apo; Magenta= NT2+KRAS. BRAF residues 1-152 (BSR) = peptides 409-560. BRAF residues 153-224 (RBD) = peptides 561-632. Red boxes highlight decreased H-D exchange rate.
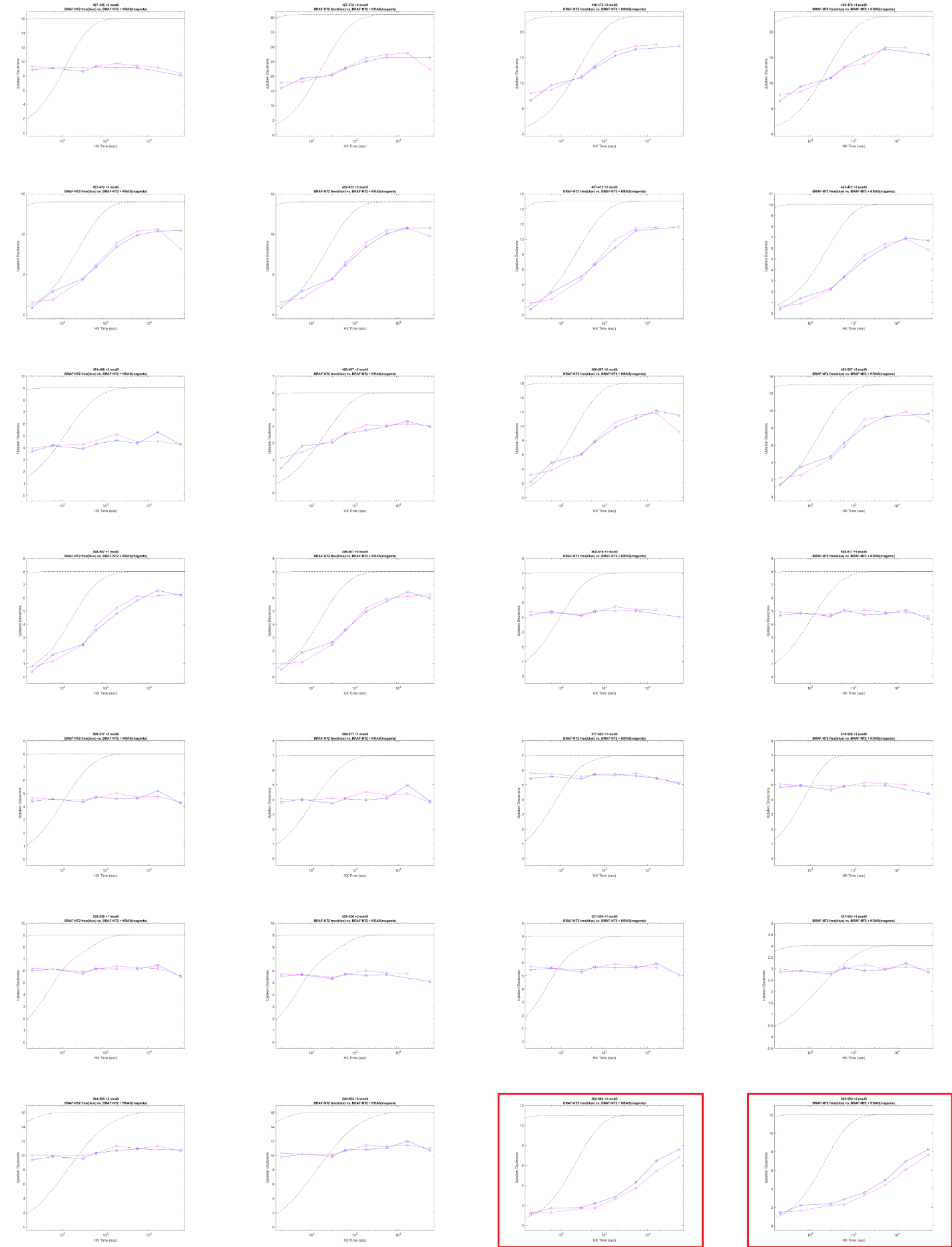

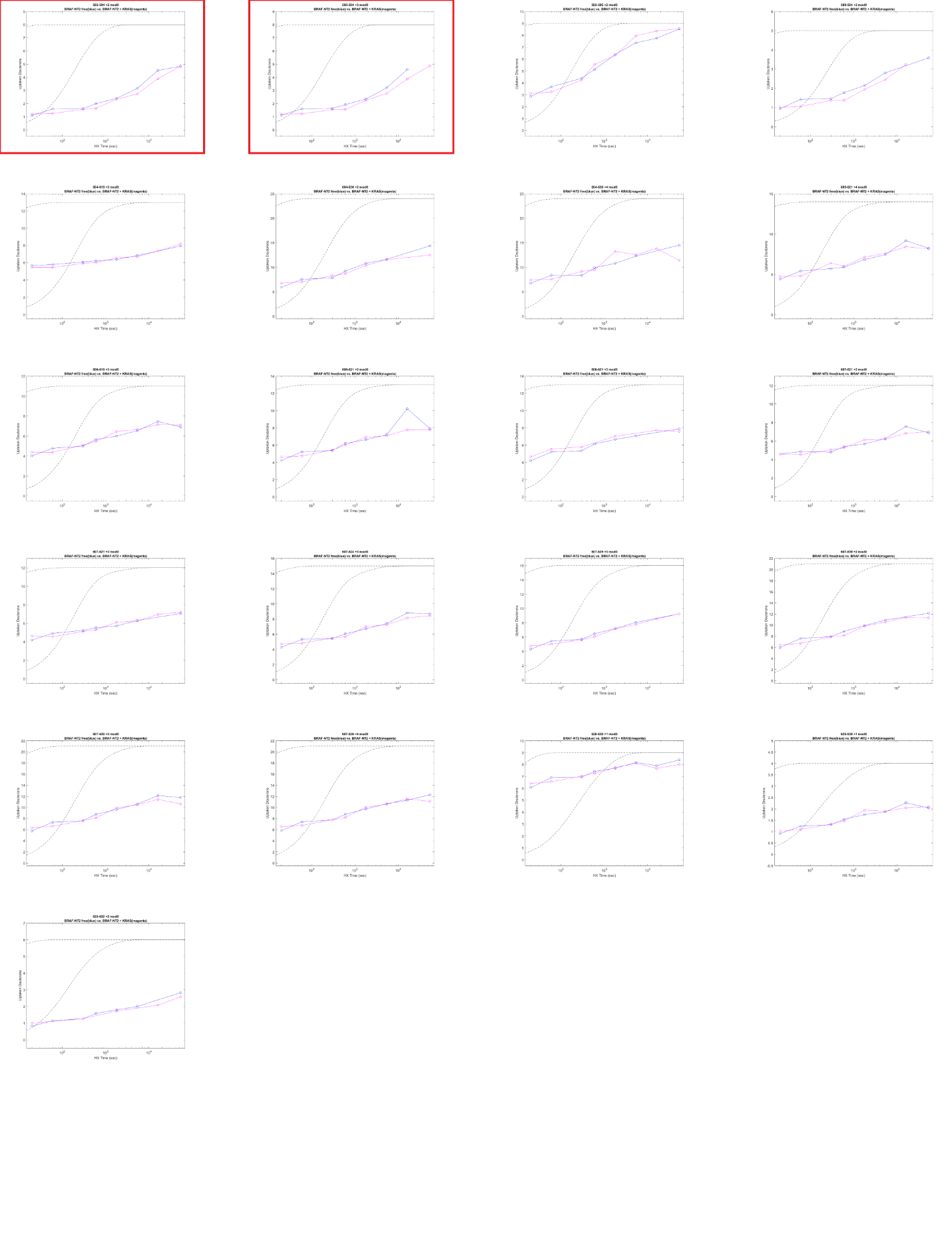
